# Quantitative tests of albendazole resistance in beta-tubulin mutants

**DOI:** 10.1101/2024.04.11.589070

**Authors:** J.B. Collins, Skyler A. Stone, Emily J. Koury, Anna G. Paredes, Fiona Shao, Crystal Lovato, Michael Chen, Richelle Shi, Anwyn Y. Li, Isa Candal, Khadija Al Moutaa, Nicolas Moya, Erik C. Andersen

**Author notes:** **Corresponding author:** Erik C. Andersen, Department of Biology, Johns Hopkins University, Bascom UTL 383, 3400 North Charles St., Baltimore, MD 21218. Equal contribution (J.B. Collins, Skyler A. Stone).

## Abstract

Benzimidazole (BZ) anthelmintics are among the most important treatments for parasitic nematode infections in the developing world. Widespread BZ resistance in veterinary parasites and emerging resistance in human parasites raise major concerns for the continued use of BZs. Knowledge of the mechanisms of resistance is necessary to make informed treatment decisions and circumvent resistance. Benzimidazole resistance has traditionally been associated with mutations and natural variants in the *C. elegans* beta-tubulin gene *ben-1* and orthologs in parasitic species. However, variants in *ben-1* alone do not explain the differences in BZ responses across parasite populations. Here, we examine the roles of five *C. elegans* beta-tubulin genes (*tbb-1*, *mec-7*, *tbb-4*, *ben-1*, and *tbb-6*) to identify the role each gene plays in BZ response. We generated *C. elegans* strains with a loss of each beta-tubulin gene, as well as strains with a loss of *tbb-1*, *mec-7*, *tbb-4*, or *tbb-6* in a genetic background that also lacks *ben-1* to test beta-tubulin redundancy in BZ response. We found that only the individual loss of *ben-1* conferred a substantial level of BZ resistance, although the loss of *tbb-1* was found to confer a small benefit in the presence of albendazole (ABZ). The loss of *ben-1* was found to confer an almost complete rescue of animal development in the presence of 30 µM ABZ, likely explaining why no additive effects caused by the loss of a second beta-tubulin were observed. We demonstrate that *ben-1* is the only beta-tubulin gene in *C. elegans* where loss confers substantial BZ resistance.

**Highlights:**

- Loss of *ben-1* provides almost complete rescue of development in albendazole (ABZ)
- Loss of different beta-tubulin genes does not confer ABZ resistance
- Loss of *ben-1* and a second beta-tubulin does not enhance the *ben-1* level of ABZ resistance

## 1. Introduction

Parasitic nematode infections are among the most common infectious diseases of humans and pose significant health and socioeconomic risks for endemic regions. Upwards of 1.5 billion individuals are estimated to be infected with at least one parasitic nematode species globally, with infections causing anemia, impaired cognitive development, reduced growth, diarrheal disease, intestinal obstructions, and lymph edema (Salikin et al., 2020). Anti-helminth drugs, or anthelmintics, are used in endemic areas to control infections and limit adverse health effects caused by parasitic nematodes. Anthelmintics are often delivered through mass drug administration (MDA) programs designed to deliver essential medicines to regions with infected populations.

One of the most common anthelmintics delivered in MDA programs is albendazole (ABZ), a drug belonging to the benzimidazole (BZ) class of anthelmintics. The BZ drug class is included in many MDA programs because of its broad-spectrum activity, capable of treating a wide variety of intestinal helminths, as well as being safe and affordable to easily deliver to large populations (Banerjee et al., 2023). Studies of the mode of BZ action have found that they inhibit the polymerization of microtubules by targeting beta-tubulin (Hastie and Georgopoulos, 1971; Sheir-Neiss et al., 1978). A study of BZ response in the free-living model nematode *Caenorhabditis elegans* found that larvae exposed to BZs were developmentally impaired and uncoordinated in locomotion (Chalfie and Thomson, 1982). Subsequent experiments showed that animals with loss-of-function mutations in the beta-tubulin gene *ben-1* were found to exhibit wild-type growth and movement in the presence of BZs (Driscoll et al., 1989). Wild-type growth, despite the loss of *ben-1*, is thought to be possible because another beta-tubulin gene acts redundantly and compensates for the loss of *ben-1*. The *C. elegans* genome contains five additional beta-tubulin genes (*tbb-1, tbb-2, mec-7, tbb-4,* and *tbb-6)* that are differentially expressed in various tissues and are thought to supply beta-tubulin function when *ben-1* is lost (Hurd, 2018).

Orthologs of *ben-1* were found to be the target of BZs in parasitic nematodes. A beta-tubulin gene (*tbb-isotype-1*) from *Haemonchus* co*ntortus*, a small-ruminant parasite, was found to rescue BZ susceptibility when expressed in a *C. elegans* strain that lacked *ben-1* (Kwa et al., 1995, 1994, 1993). Unlike *C. elegans,* the *H. contortus* genome contains only four genes encoding beta-tubulins (*tbb-isotype-1*, *tbb-isotype-2*, *tbb-isotype-3*, and *tbb-isotype-4*). A smaller complement of beta-tubulin genes, combined with expression differences between each of the four genes has led to the conclusion that loss of *tbb-isotype-1* likely causes lethality, indicating that BZ resistance in parasites is probably dependent on altered function variants in beta-tubulin. However, parasitic nematodes currently lack the genetic tools, such as genome editing, to validate resistance genes using targeted mutations. Exploration of anthelmintic resistance is dependent on *C. elegans* as a complement to research in parasites, and a cycle of discovery has been proposed to explore and validate the mechanisms of BZ resistance using both free-living and parasitic nematodes (Wit et al., 2021).

Anthelmintic resistance is a major concern in the control of parasites. Resistance to the BZ drug class has become nearly ubiquitous in many nematode species of veterinary importance and is now an emerging problem in nematode infections of humans (Howell et al., 2008; Kaplan, 2004; Krücken et al., 2017). The development of resistance to BZs makes the control of infections difficult and costly. To address the emergence of BZ resistance, it is necessary to understand the underlying genetics contributing to resistance. After suspected resistance-associated variants are identified in parasites, they can be validated in *C. elegans* using CRISPR-Cas9 genome editing. Studies of BZ resistance have identified non-synonymous variants at codons 134, 167, 198, and 200 of *ben-1* orthologs in parasites (Avramenko et al., 2019; Kwa et al., 1994; Mohammedsalih et al., 2020; Venkatesan et al., 2023). Every known beta-tubulin variant associated with BZ resistance in parasitic nematodes has been shown to cause resistance in *C. elegans* by the introduction of the variant into the *ben-1* gene (Dilks et al., 2021, 2020; Kitchen et al., 2019; Kwa et al., 1994; Venkatesan et al., 2023). These variants in parasite beta-tubulin genes are thought to alter a putative BZ binding site, preventing BZs from inhibiting beta-tubulin, preserving the normal formation of microtubules, and allowing nematodes to survive and develop normally in the presence of BZ treatment.

Despite the validation of variants in *ben-1* orthologs as a mechanism of resistance to BZs, *ben-1* is not the only gene involved in BZ resistance. Genome-wide association studies in wild populations of *C. elegans* have identified multiple genomic loci independent of *ben-1* that are associated with BZ resistance (Hahnel et al., 2018; Zamanian et al., 2018). Fully understanding the genetics of resistance is necessary to inform strategic decisions that improve the efficacy of existing treatments, as well as lead to the development of new treatments and control strategies. Thus, it is imperative to identify all genes associated with BZ resistance. Here, we explore the effects that loss of each beta-tubulin gene has on BZ resistance in *C. elegans*. The gene *ben-1* has been extensively studied and confers the greatest level of BZ resistance. However, the roles of the other *C. elegans* beta-tubulin genes (*tbb-1*, *tbb-2*, *mec-7*, *tbb-4*, and *tbb-6*) in BZ resistance are not well understood. We have compared the effects of single gene deletions of each beta-tubulin gene on nematode development when exposed to a single concentration of ABZ that previously has been found to confer a significant impact on the development of the wild-type N2 strain of *C. elegans* (Dilks et al., 2021, 2020). We find that the loss of *ben-1* conferred the highest level of resistance and the loss of *tbb-1* conferred moderate resistance. To test for genetic redundancy among beta-tubulin genes, we used CRISPR-Cas9 genome editing to delete each beta-tubulin gene in a genetic background that already has lost *ben-1* function. The loss of each beta-tubulin gene in the *ben-1* deletion background did not confer a detectable change in ABZ resistance compared to the loss of *ben-1* alone. Overall, we find that the loss of *ben-1* alone is sufficient to confer the maximum level of *C. elegans* ABZ resistance at the concentration tested.

## 2. Materials and Methods

### 2.1 Generation of phylogeny of selected nematode beta-tubulins

Five nematode species were selected to make a phylogenetic tree of beta-tubulins to observe levels of conservation. All nematode species selected are Clade V nematodes as the association of *ben-1* orthologs with BZ resistance has most often been validated in this clade. *C. elegans* and *Caenorhabditis briggsae* were selected as two closely related free-living nematode species. *Pristionchus pacificus*, another free-living nematode, was selected because of its high-quality genome and evolutionary divergence from *C. elegans*. Many parasite genomes are relatively poor quality and lack detailed gene annotations, so we chose two parasite species with well annotated genomes, the hookworm *Necator americanus* and *H. contortus*, to include in the phylogenetic tree.

*Orthofinder* (Emms and Kelly, 2019) was used to identify beta-tubulin sequences (Supplementary Table 1) from each species. Data were obtained from the following sources: WormBase Parasite (WBPS18) (*H. contortus*, *N. americanus*, *P. pacificus*), WormBase (WS279) (*C. elegans*), and from a previous publication (*C. briggsae*) (Moya et al., 2023). Ortholog sequences were aligned using *Mafft,* and the phylogenetic tree was generated and annotated using *IQTREE* (Katoh et al., 2002; Minh et al., 2020). *IQTREE* performs automatic model selection. The selected model was LG+G4, which uses the LG model (Le and Gascuel, 2008) to examine amino-acid exchange rates and a discrete gamma model with four categories (G4) (Yang, 1994) to examine heterogeneity across amino acid sites. Branch support was estimated with 1000 iterations of ultrafast bootstrap approximation (Minh et al., 2013). Putative clades were identified in the generated tree and colored by clade.

### 2.2 C. elegans strains and maintenance

Nematodes were grown on plates of modified nematode growth media (NGMA) containing 1% agar and 0.7% agarose and seeded with the *Escherichia coli* strain OP50 (Andersen et al., 2014). Plates were maintained at 20°C for the duration of all experiments. Before each assay, animals were grown for three generations to reduce the multigenerational effects of starvation.

CRISPR-Cas9-edited strains were generated as previously described (Dilks et al., 2020; Hahnel et al., 2018) (Supplementary File 2), except for VC364 *tbb-1(gk207)*, which was acquired from the *Caenorhabditis* Genetics Center (Minneapolis, MN). All single deletions were generated in the reference N2 genetic background. All double deletions were generated in the ECA882 *ben-1(ean64)* genetic background (Dilks et al., 2021, 2020). Progeny from injected animals (F1) were individually placed onto NGMA plates to reproduce and then sequenced using Sanger sequencing to confirm the presence of the desired edit. At least two generations of animals after single-animal passage were Sanger sequenced to confirm successful genome edits. Two independent edits of each strain were generated to control for any potential off-target effects caused by CRISPR-Cas9.

### 2.3 Nematode food preparation

The OP50 strain of *E. coli* was used as a nematode food source on NGMA plates. Bacterial food for the liquid-based high-throughput assay was prepared as previously described (Widmayer et al., 2022). Briefly, a frozen stock of the HB101 strain of *E. coli* was used to inoculate and grow a one liter culture at an OD_600_ value of 0.001. Six cultures containing one liter of pre-warmed 1x Horvitz Super Broth (HSB) and an OD_600_ inoculum grew for 15 hours at 37°C until cultures were in the late log growth phase. After 15 hours, flasks were removed from the incubator and transferred to 4°C to halt bacterial growth. Cultures were pelleted using centrifugation, the supernatant removed, and washed with K medium. Bacteria were resuspended in K medium, and the OD_600_ value was determined. The bacterial suspension was diluted to a final concentration of OD_600_100 before being aliquoted to 30 mL and frozen at -80°C.

### 2.4 Albendazole stock preparation

A 100 µM stock solution of albendazole (Fluka, Catalog #: A4673-10G) was prepared in dimethyl sulfoxide (DMSO), aliquoted, and stored at -20°C. A frozen ABZ aliquot was defrosted shortly before adding the drug to the assay plates.

### 2.5 High-throughput phenotyping assay (HTA)

A previously described HTA was used for all ABZ response phenotyping assays (Shaver et al., 2023). Two independent assays made up of three bleaches each were performed. Strains underwent three generations of growth to control for any starvation effects and were then bleach synchronized in triplicate to control for variation caused by bleach effects. Embryos were concentrated at 0.6 embryos/µL in 50 µL of K medium (Boyd et al., 2012). A volume of 50 µL of the embryo solution was dispensed into each well of a 96-well plate. Both DMSO and ABZ conditions contained 48 wells of N2 and ECA882, and 24 wells of each of the other tested strains for each replicate bleach. Embryos were allowed to hatch overnight at 20°C with constant shaking at 180 rpm. The following morning, HB101 aliquots were thawed at room temperature, combined, and diluted to OD_600_30 with K medium, and kanamycin was added at a concentration of 150 µM to inhibit further bacterial growth and prevent contamination. The final well concentration of HB101 was OD_600_10 and the final concentration of kanamycin was 50 µM, and each well was treated with either 1% DMSO or 30 µM ABZ in 1% DMSO. Animals were grown for 48 hours with constant shaking at 180 rpm, after which, animals were treated with 50 mM sodium azide in M9 buffer to straighten and paralyze the animals for imaging. Following 10 minutes of exposure to sodium azide, each plate was imaged using a Molecular Devices ImageXpress Nano microscope (Molecular Devices, San Jose, CA) with a 2X objective (Shaver et al., 2023).

Independent assays included identical strain sets except as follows: Strains with a deletion of *tbb-2* were found to be too developmentally delayed to use in these assays. The ECA3746 *ben-1(ean64)*; *mec-7(ean257)* strain was removed from assay one because of an insufficient quantity of embryos after bleach synchronization. Smaller significant effects on animal development were observed for some single deletions in control conditions of assay one but not in assay two, indicating that significance assigned to the observed small effects could be the result of high levels of replication, making even small differences significant.

### 2.6 Data cleaning and analysis

High-throughput assay images were processed using CellProfiler (https://github.com/AndersenLab/CellProfiler). Processed image data were cleaned and processed using the *easyXpress* (Nyaanga et al., 2021) R package as previously described (Shaver et al., 2024). The two assays were cleaned and processed independently. All statistical comparisons and figure generation were performed in R(4.1.2) (R Core Team, 2020). We used the *Rstatix* package *tukeyHSD* function on an ANOVA model generated with the formula *phenotype ∼ strain* to calculate differences in the responses of the strains. Figure 3 was generated using data from assay one because of the large amount of variation shown in animal response for the VC364 *tbb-1(gk207)* strain in assay two, thought to be caused by human error. Figure 4 was generated using data from assay two, because of the loss of the ECA3746 strain in assay one. All data are presented in supplemental figures.

## 3. Results

### 3.1 Beta-tubulins are well conserved among Clade V nematode species

We wanted to determine how each of the six beta-tubulin genes from *C. elegans* were related to each other, as well as to orthologs from other nematode species (Hurd, 2018). Phylogenetic analysis found five putative clades of beta-tubulin proteins (Figure 1). *Caenorhabditis elegans tbb-1* and *tbb-2* share a common clade with the *tbb-isotype-1* beta-tubulins from *H. contortus* and *N. americanus*. *Caenorhabditis elegans mec-7* and *tbb-4* are in separate clades with *tbb-isotype-3* and *tbb-isotype-4* clustering with each gene, respectively. The genes *ben-1* and *tbb-isotype-2* each cluster into separate clades. The gene *tbb-6*, a beta-tubulin unique to *C. elegans,* could not be placed into the tree because of a high level of divergence. The high levels of conservation of beta-tubulins among Clade V species highlight the ability to use *C. elegans* as a model system to investigate the broad roles of beta-tubulins in BZ resistance across diverse nematode species.

**Figure 1.**
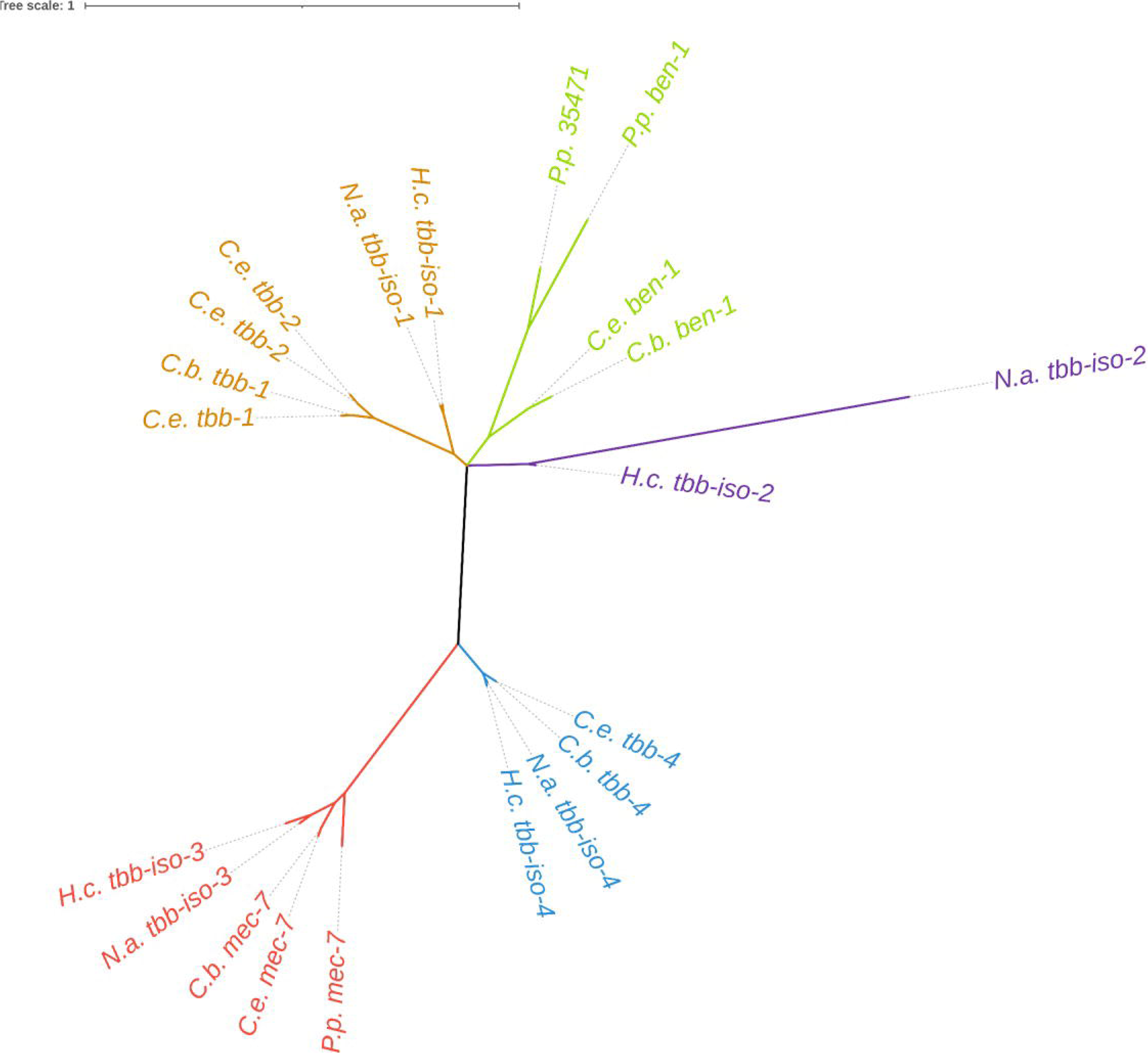
Phylogenetic relationship of nematode beta-tubulins. Beta-tubulin gene models from three free-living nematodes, *Caenorhabditis elegans* (*C.e.*), *C. briggsae* (*C.b.*), and *Pristionchus pacificus* (*P.p.*), and two parasitic nematodes, *Haemonchus contortus* (*H.c.*) and *Necator americanus* (*N.a.*), were used to generate a tree showing the relationship between beta-tubulin genes. Branches are colored by putatively assigned clades. Sequence data were obtained from the following sources: WormBase Parasite (WBPS18) (*H. contortus*, *N. americanus*, *P. pacificus*), WormBase (WS279) (*C. elegans*), and from a previous publication (*C. briggsae*) (Moya et al. 2023).

### 3.2 The loss of ben-1 is the only beta-tubulin gene to confer high levels of ABZ resistance

CRISPR-Cas9 genome editing was used to generate deletions of each beta-tubulin gene in the N2 laboratory strain genetic background (Figure 2). Edited strains with single deletions of each beta-tubulin gene were phenotyped in DMSO and ABZ using a previously described high-throughput assay (HTA) that quantitatively measures nematode development (Shaver et al., 2023; Widmayer et al., 2022). Briefly, strains were bleach synchronized and embryos were titered into 96-well plates. The following day, arrested L1 larvae were given OP50 *E. coli* with either 1% DMSO or 30 µM ABZ and 1% DMSO. Plates were incubated for 48 hours at 20°C with constant shaking at 180 rpm. Animals were then treated with sodium azide and imaged to quantify the lengths of each animal in each well of a 96-well plate. Median animal lengths were calculated from each well of an assay plate and normalized across independent growths, plates, and bleaches. Deletion of each beta-tubulin gene in the same genetic background enables the determination of the quantitative effects that each gene has on BZ response, as well as to determine if the loss of each beta-tubulin gene impacts development in control conditions. Median animal length after 48 hours of exposure was normalized to control conditions, and then statistical comparisons were made between N2 and each strain. The loss of *tbb-1* had the most significant impact on development in control conditions, indicating that the loss of *tbb-1* is detrimental (S Figs. 4,6). The loss of *ben-1* was the only strain to confer high levels of resistance to ABZ, almost fully rescuing development compared to control conditions (Figure 3, S Figs. 3,5). The loss of *tbb-1* was found to confer a moderate level of resistance, with animal development significantly less affected than the wild-type strain but still heavily affected by ABZ as compared to control conditions.

**Figure 2.**
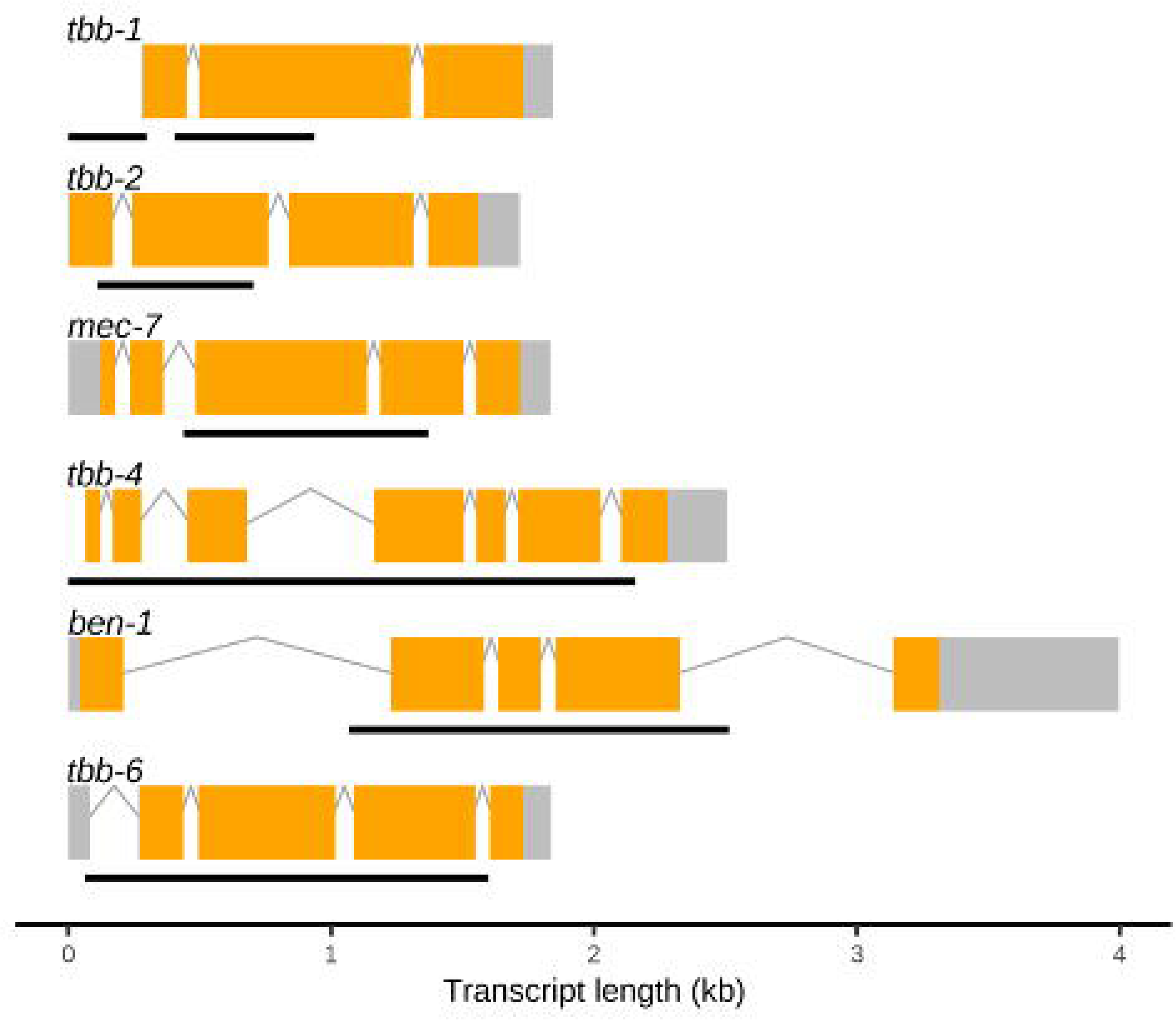
Gene models and locations of deletion alleles generated in *C. elegans* beta-tubulin genes. Gene models of the longest isoforms are presented for each *C. elegans* beta-tubulin gene, with exons (orange), introns (gray lines), and untranscribed regions (gray boxes) shown. Regions that were deleted using CRISPR-Cas9 genome editing are shown as black lines under each model. Deleted regions of *tbb-1* are shown as two black lines because strains with two independent deletion alleles were used. Gene model data were obtained from WormBase (WS279).

**Figure 3.**
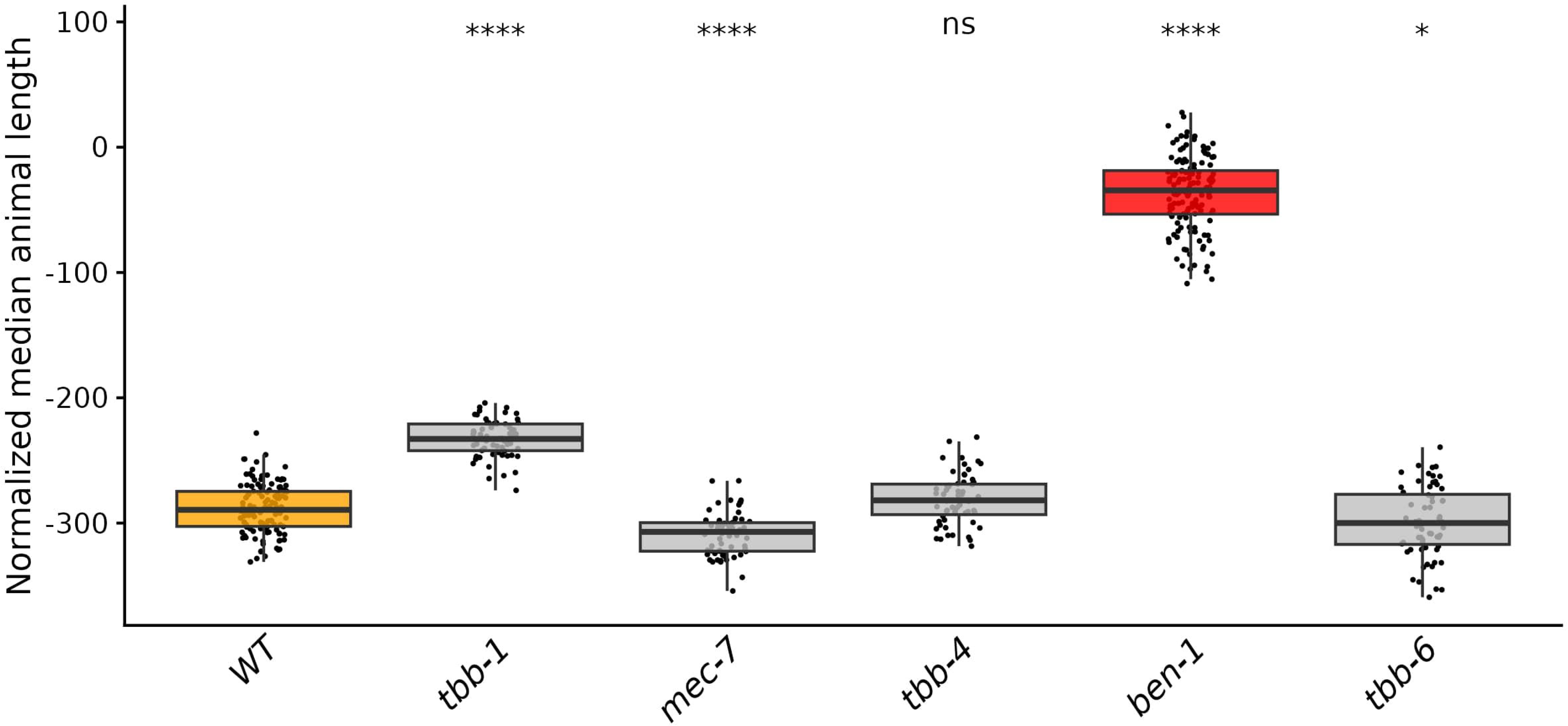
Only loss of *ben-1* causes resistance to ABZ. Median animal lengths of strains grown in 30 µM ABZ that have been regressed for bleach effects and then normalized to the mean of all median animal lengths from the control condition are shown. Each point represents the summarized measurements of an individual well containing five to 30 animals. Data are shown as box plots with the median as a solid horizontal line and the 75th and 25th quartiles on the top and bottom of the box, respectively. The top and bottom whiskers extend to the maximum point within the 1.5 interquartile range from the 75th and 25th quartiles, respectively. Statistical significance compared to the wild-type strain is shown above each strain (*p* < 0.05 = *, *p* < 0.0001 = ****, ANOVA with Tukey HSD).

**Figure 4.**
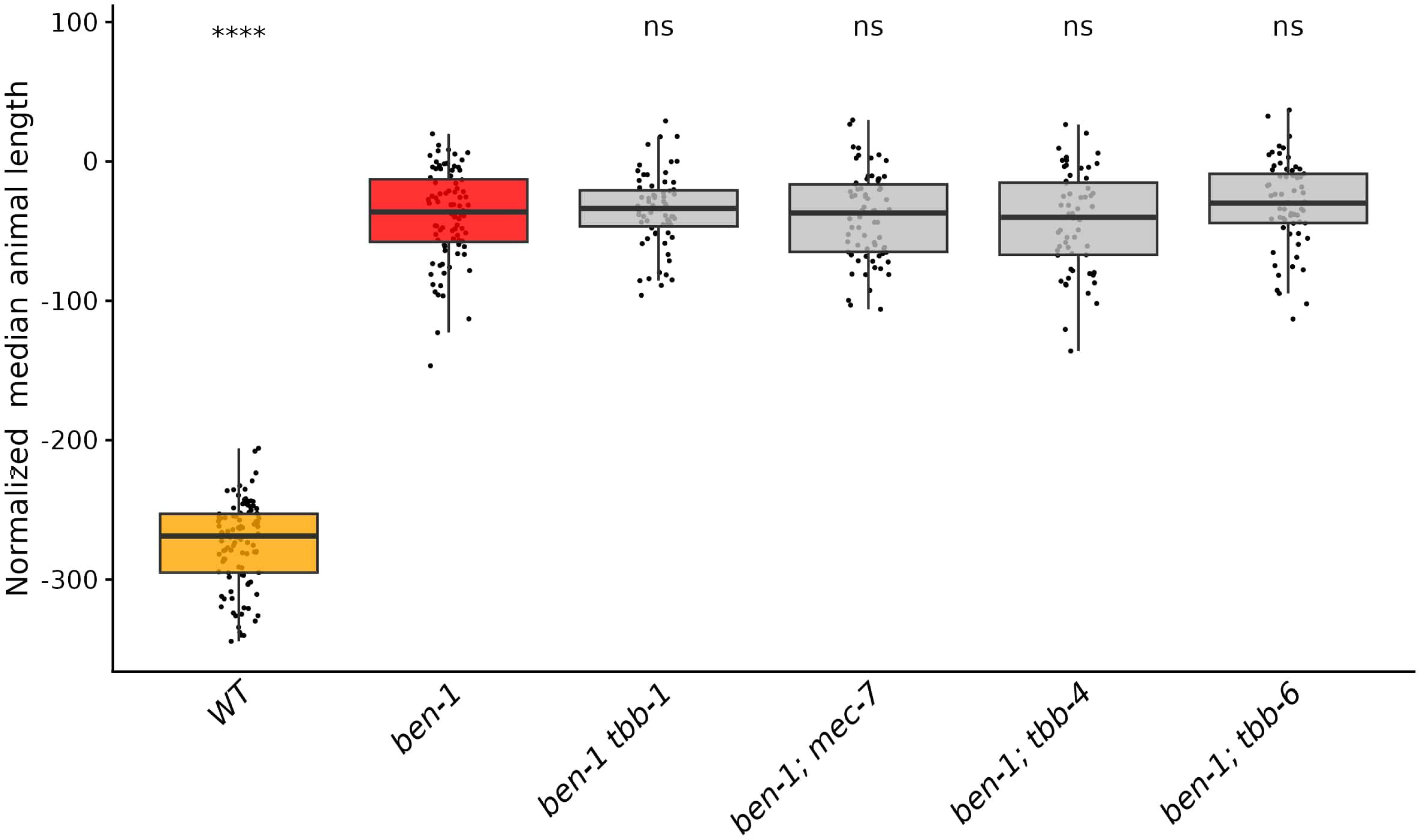
None of the other beta-tubulin genes act redundantly with *ben-1* in ABZ response. Median animal lengths of strains grown in 30 µM ABZ that have been regressed for bleach effects and then normalized to the mean of all median animal lengths from the control condition are shown. Each point represents the summarized measurements of an individual well containing five to 30 animals. Data are shown as box plots with the median as a solid horizontal line and the 75th and 25th quartiles on the top and bottom of the box, respectively. The top and bottom whiskers extend to the maximum point within the 1.5 interquartile range from the 75th and 25th quartiles, respectively. Statistical significance compared to the Δ*ben-1* strain is shown above each strain (*p* < 0.05 = *, *p* < 0.0001 = ****, ANOVA with Tukey HSD).

### 3.3 The loss of ben-1 confers the highest level of ABZ resistance compared to other beta-tubulin mutants

To determine if other beta-tubulin genes play a redundant role in ABZ resistance with *ben-1*, we generated individual deletions of *tbb-1*, *mec-7*, *tbb-4*, and *tbb-6* in the *ben-1(ean64)* genetic background. We exposed these double beta-tubulin mutants to DMSO and ABZ in the same high-throughput development assay described above to determine if the loss of a second beta-tubulin alters the levels of BZ resistance observed in the single *ben-1* mutant. Similarly to the single deletion assay, small significant differences were observed for multiple strains compared to the wild-type strain in control conditions, except for the strain ECA3628 *ben-1(ean64)*; *tbb-4(ean282)* (S. Figs 8,10), which likely has off-target effects of gene editing that impacted growth compared to the independently edited second strain. Small differences in the summarized median length reflect differences in the developmental rate that could be caused by the combined effects of the loss of multiple beta-tubulins. Strains with the loss of a second beta-tubulin were found to be equally resistant when compared to the loss of *ben-1* alone (Figure 4, S Figs. 7,9). As previously noted, the loss of *ben-1* almost fully rescued development at 30 µM ABZ compared to the control strain, possibly preventing any small effects conferred by the loss of a second beta-tubulin from being observed.

## 4. Discussion

Despite the role beta-tubulin variants have in BZ resistance, the collective understanding of BZ resistance comes from studies of *C. elegans ben-1* and orthologs in parasites. Fully understanding the mechanisms underlying BZ resistance is imperative to the future of BZs as anthelmintic treatments. Here, we take an important first step to test additional beta-tubulin genes in BZ resistance.

### 4.1 ben-1 plays the largest role in ABZ resistance in C. elegans

We examined the role that five of the six *C. elegans* beta-tubulin genes play in ABZ resistance by generating strains with a loss of each gene, as well as strains with a loss of an additional beta-tubulin in a *ben-1* mutant background. Because of detrimental effects on development, strains with a loss of *tbb-2* could not be measured for responses to ABZ. Consistent with previous studies, the loss of *ben-1* was sufficient to confer the maximum level of ABZ resistance, though it is important to note that the loss of *tbb-1* conferred a moderate level of resistance. Loss of a second beta-tubulin in a strain with a loss of *ben-1* did not confer a detectable enhancement of resistance. However, we can not definitively conclude if any other beta-tubulin gene acts redundantly with *ben-1* in ABZ resistance. The assay that we used to measure ABZ resistance uses one concentration that previously was found to differentiate susceptible strains from *ben-1* mutant strains (Dilks et al., 2021, 2020). It remains possible that enhancement of ABZ resistance could be detected at higher ABZ concentrations where the single contribution of *ben-1* might not be sufficient to cause resistance alone. Another caveat is that only a single trait, development, was measured. ABZ affects multiple traits, including fecundity and competitive fitness over multiple generations (Shaver et al., 2024). Future studies should investigate multiple traits at different ABZ concentrations to fully understand the role of all beta-tubulin genes in the ABZ response.

### 4.2 BZ resistance is complicated by differences in beta-tubulin copy number, levels of expression, and resistance alleles

We tested the role of each beta-tubulin gene in ABZ response by deleting much of the coding sequence. Therefore, these results are binary for the presence or absence of each beta-tubulin gene. Amino-acid altering variants from parasites have been validated in ABZ resistance using *C. elegans* and shown to cause ABZ resistance equivalent to a strain with a loss of *ben-1* (Dilks et al., 2021, 2020; Venkatesan et al., 2023). However, these variants likely do not cause loss of *tbb-isotype-1* function in parasites (Saunders et al., 2013). What could be causing this discrepancy between loss-of-function variants in *C. elegans* and potential altered function variants in parasitic nematodes? In species with highly expressed beta-tubulin genes that have BZ-sensitive alleles, loss-of-function alleles would cause fitness defects, similar to what we see with *tbb-1* and *tbb-2* (Figure 4). In these species, benzimidazole resistance must be mediated by altered function variants. In species with less highly expressed (or tissue-specific) beta-tubulin genes that have BZ-sensitive alleles, loss-of-function alleles could cause BZ resistance because other beta-tubulin genes can substitute for essential functions, similar to what we see with *ben-1* (Hurd, 2018). Interestingly, the *H. contortus* beta-tubulin gene *tbb-isotype-2* is shown to be equally related to *tbb-isotype-1* and *ben–1*, and loss-of-function alleles of this gene have been documented in some highly resistant *H. contortus* populations (Saunders et al., 2013). Additionally, the phenotypic classification of BZ-resistance phenotypes differs between these two species and can be explained by differences in loss-of-function vs. altered function mutations. In *C. elegans* where *ben-1* variants or mutations can cause loss of function, the BZ-resistance phenotype is recessive (Dilks et al., 2021). By contrast, putative BZ-resistance alleles in *H. contortus* are hypothesized to cause dominant BZ resistance (Silvestre et al., 2001).

Beyond coding variants or mutations in beta-tubulin genes, changes in the levels and tissue-specific expression can alter BZ resistance. Previously, we found that some *C. elegans* wild strains with clear ABZ resistance do not have variants that alter the coding sequence of *ben-1* but instead have much lower expression levels of *ben-1* as compared to the rest of the population (Zhang et al., 2022). These strains are resistant because the susceptible beta-tubulin protein is not expressed. Additionally, we found that the expression of *ben-1* in cholinergic neurons alone is sufficient to confer susceptibility to ABZ (Gibson et al., 2022), highlighting that variants modifying expression in specific tissues could confer resistance in a unique way independent of the beta-tubulin coding sequence. These observations from both *C. elegans* and *H. contortus* demonstrate that more attention should be paid to the number of beta-tubulin genes, their levels of expression, the sites of expression, and the putative BZ-resistance alleles found in each beta-tubulin gene. To definitively understand BZ resistance mediated by beta-tubulin genes, we must also drastically improve parasitic nematode genomes and gene models because most species lack full descriptions of their beta-tubulin complement.

## 5. Future directions

The role of *ben-1* and *tbb-isotype-1* beta-tubulins in BZ resistance has been thought to be similar and has established *C. elegans* as an essential model for parasite BZ resistance research. However, BZ treatment is typically fatal in susceptible parasites (Prichard, 1988), as well as documented ovicidal effects of BZs against parasite embryos (Boes et al., 1998). Conversely, the same effects are not typically seen in *C. elegans* where the most significant impact is often on the developmental rate (Shaver et al., 2022). The loss of *tbb-1* or *tbb-2* was deleterious and loss-of-function mutations in either gene would likely be rapidly selected against in the wild (*i.e.*, no variants are observed in natural *C. elegans* strains) (Crombie et al., 2024), similarly to the predicted loss of *tbb-isotype-1*. It is important to note that *tbb-1* and *tbb-2* have known resistance alleles at amino acid position 200, and future studies should edit both genes to make them harbor BZ-sensitive alleles to more closely approximate the beta-tubulin complement and alleles in *H. contortus*. Such studies could offer an improved model system for investigating BZ resistance. However, studies of BZ resistance need to investigate variants beyond single amino-acid alterations. Our results demonstrate that a variety of factors such as copy number, expression, and tissue-specific function can all affect BZ resistance. To continue to broaden our understanding of BZ resistance, we must expand to a whole-genome approach that investigates variants across every single beta-tubulin gene and beyond that single class of genes.

## Data Availability

All code and data are openly available at https://github.com/AndersenLab/2024_beta_tubulin_manuscript

## Acknowledgments

We would like to thank members of the Andersen lab for their feedback in the preparation of this manuscript. We thank the *Caenorhabditis* Natural Diversity Resource (NSF Capacity Grant 2224885) and the *Caenorhabditis* Genetics Center (NIH Office of Research Infrastructure Programs, P40 OD010440) for providing the strains used in this study. Additionally, we would also like to thank WormBase for providing an essential resource for genetic and genomic data used in this manuscript.

---

**Supplemental Figure 1.**
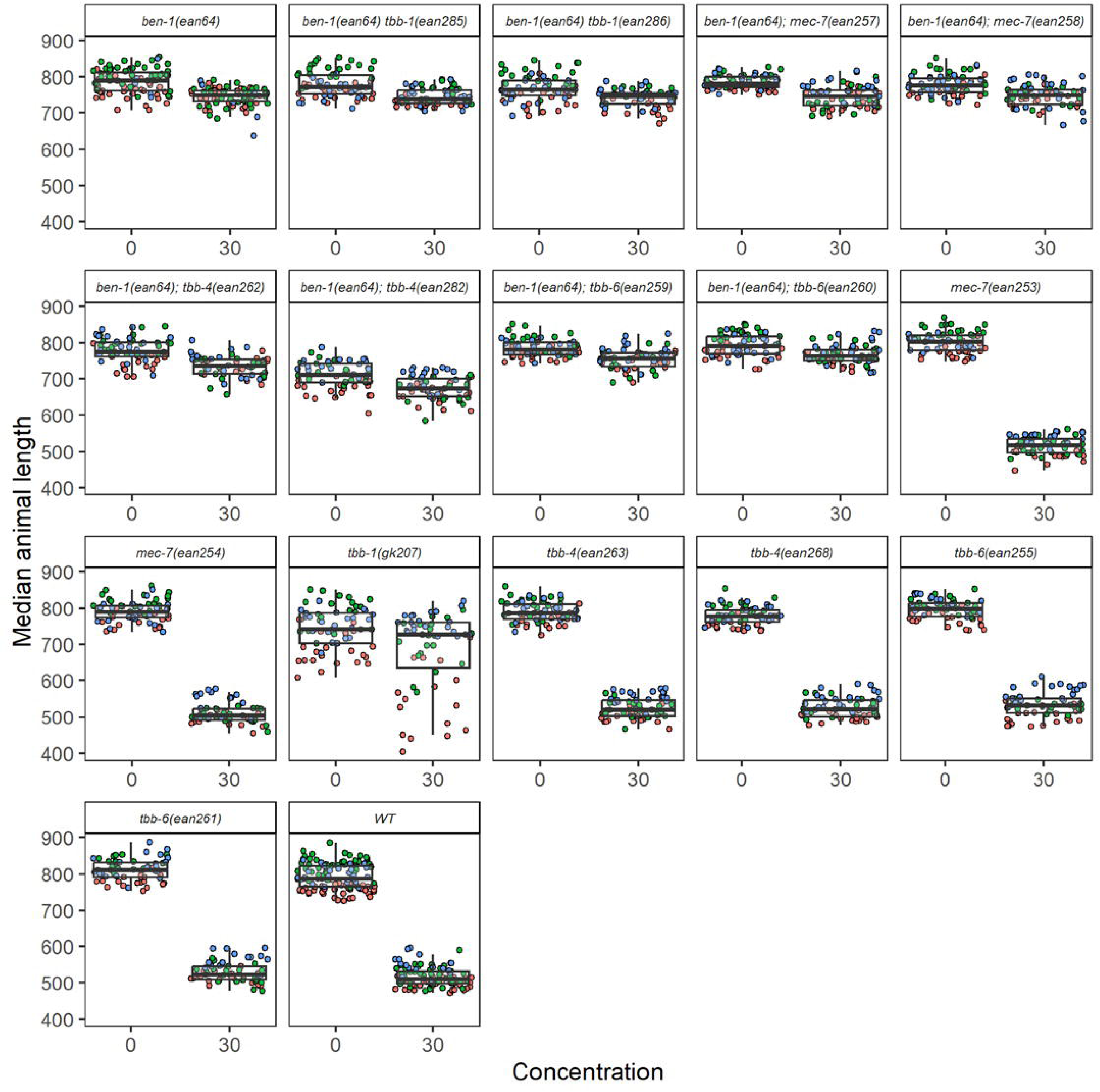
Distribution of raw animal lengths for each strain in assay two after exposure to ABZ. Raw median animal lengths, summarized by well, for each strain are shown for DMSO (0 µM ) and ABZ (30 µM) conditions. Wells are colored by the corresponding replicate bleach synchronization (red=1, green=2, blue=3).

**Supplemental Figure 2.**
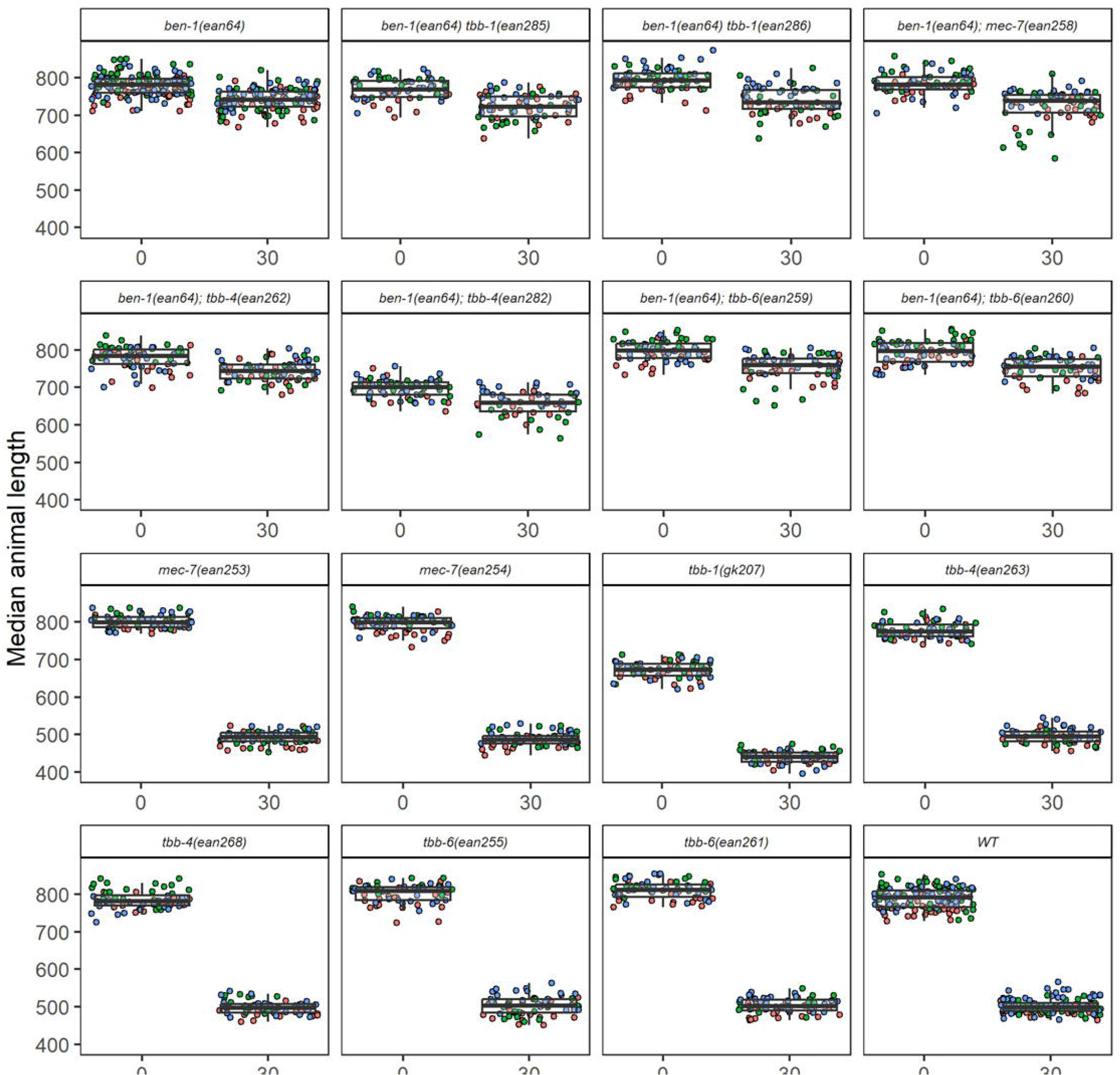
Distribution of raw animal lengths for each strain in assay one after exposure to ABZ. Raw median animal lengths, summarized by well, for each strain are shown for DMSO (0 µM ) and ABZ (30 µM) conditions. Wells are colored by the corresponding replicate bleach synchronization (red=1, green=2, blue=3).

**Supplemental Figure 3.**
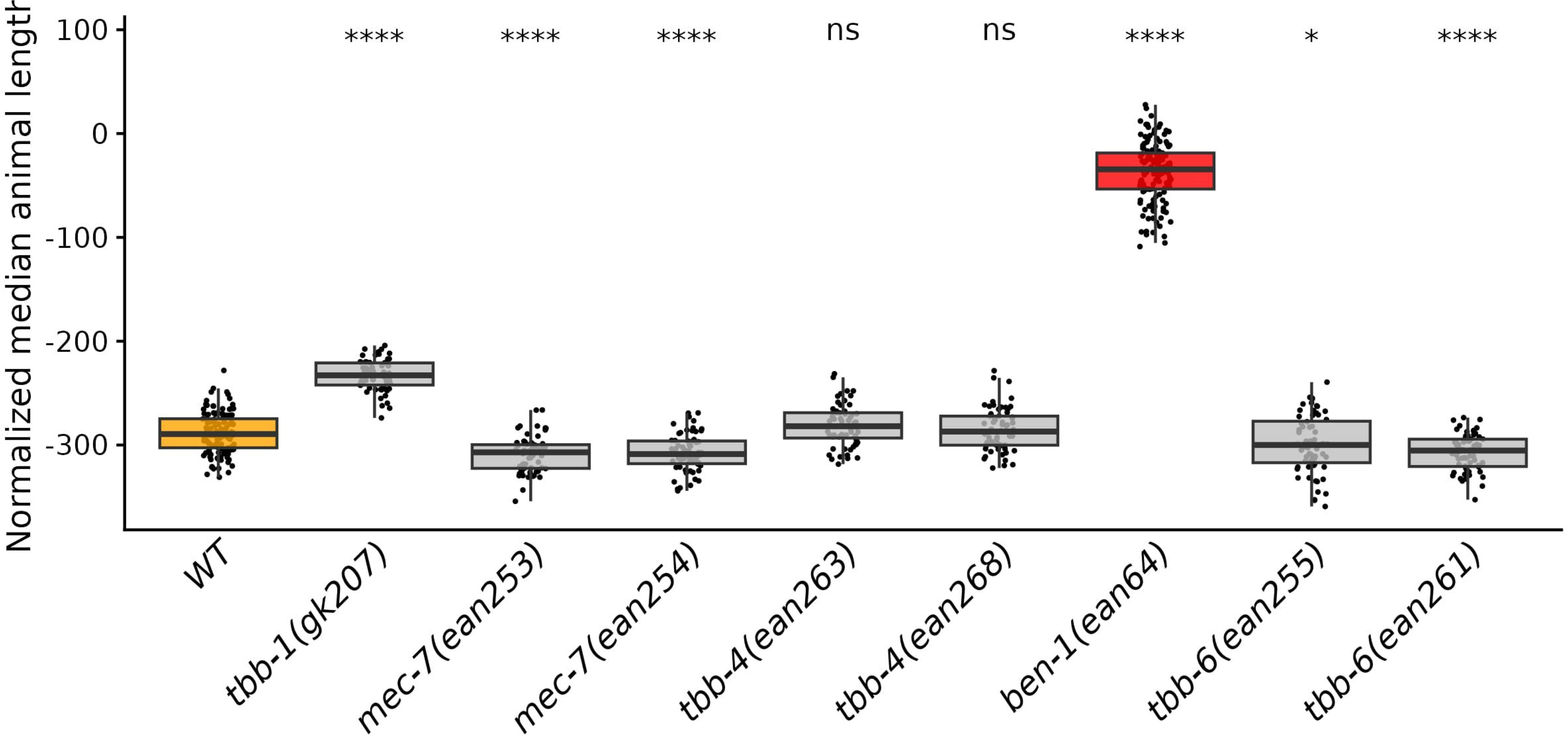
Only loss of *ben-1* causes ABZ resistance. Median animal lengths of strains grown in 30 µM ABZ that have been regressed for bleach effects and then normalized to the mean of all median animal lengths from the control condition are shown. Each point represents the summarized measurements of an individual well containing five to 30 animals. Data are shown as box plots with the median as a solid horizontal line and the 75th and 25th quartiles on the top and bottom of the box, respectively. The top and bottom whiskers extend to the maximum point within the 1.5 interquartile range from the 75th and 25th quartiles, respectively.. Statistical significance compared to the wild-type strain is shown above each strain (*p* < 0.05 = *, *p* < 0.0001 = ****, ANOVA with Tukey HSD).

**Supplemental Figure 4.**
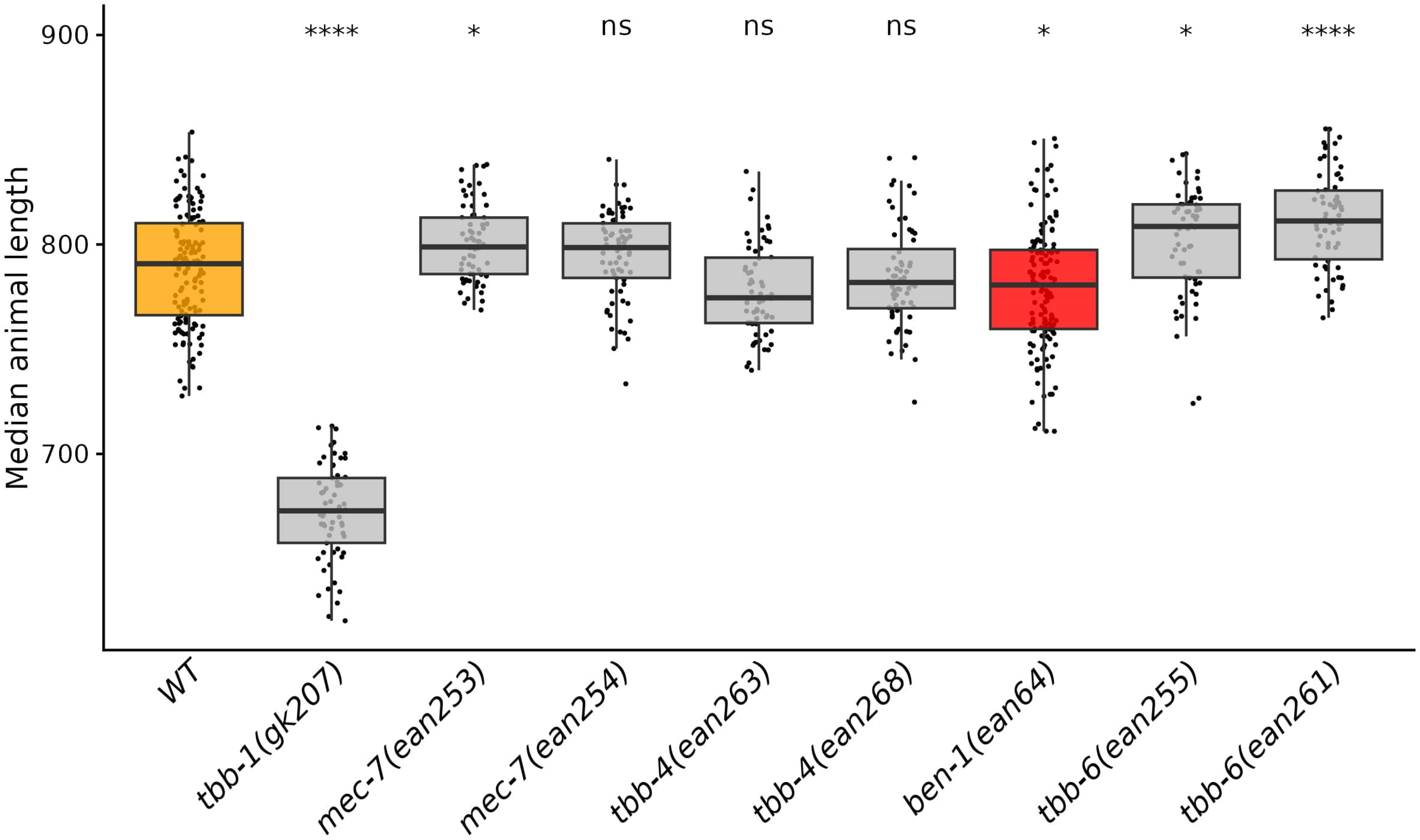
Loss of beta-tubulin genes affects animal lengths in control conditions. Median animal lengths of strains grown in 1% DMSO are shown. Each point represents the summarized measurements of an individual well containing five to 30 animals. Data are shown as box plots with the median as a solid horizontal line and the 75th and 25th quartiles on the top and bottom of the box, respectively. The top and bottom whiskers extend to the maximum point within the 1.5 interquartile range from the 75th and 25th quartiles, respectively. Statistical significance compared to the wild-type strain is shown above each strain (*p* < 0.05 = *, *p* < 0.0001 = ****, ANOVA with Tukey HSD).

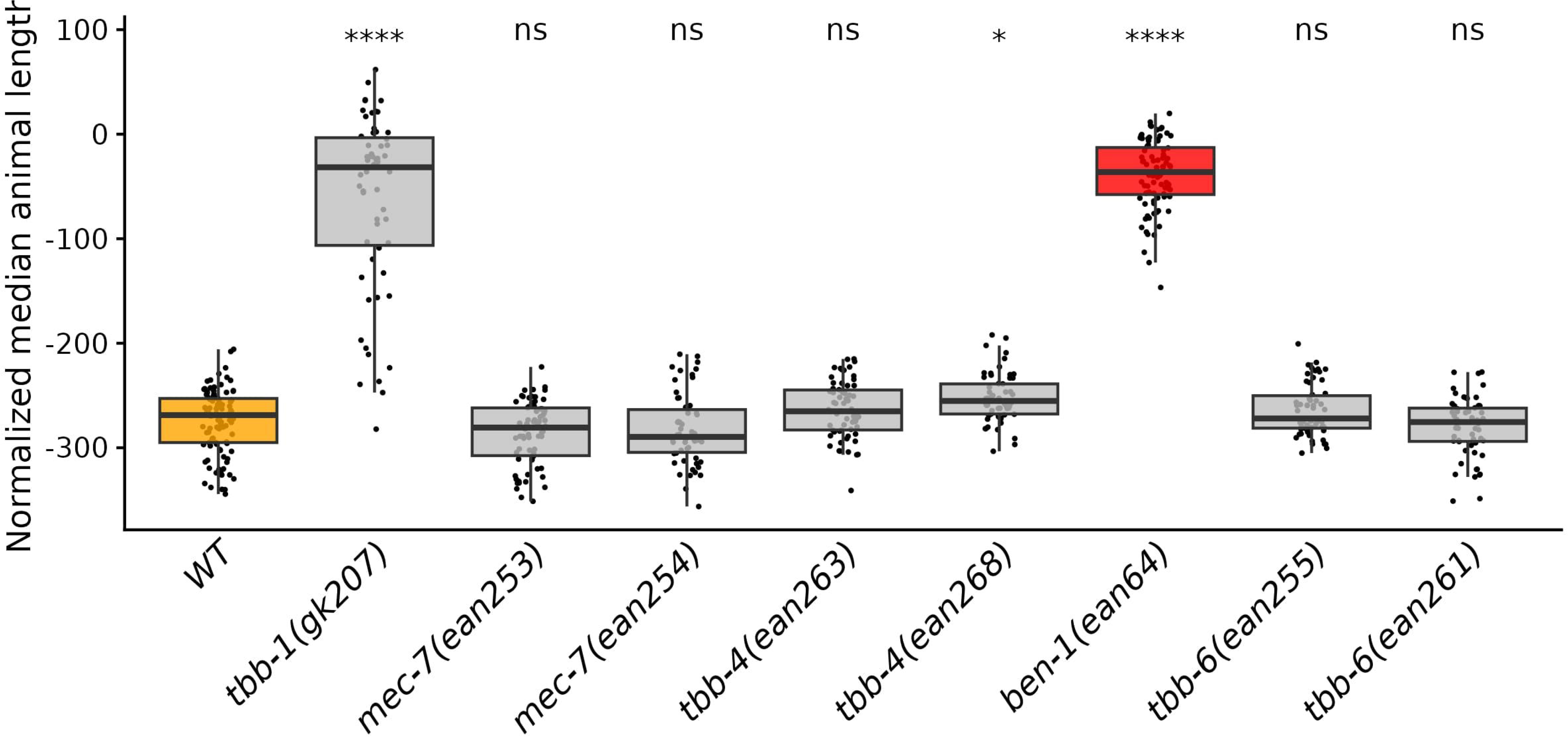

**Supplemental Figure 6.**
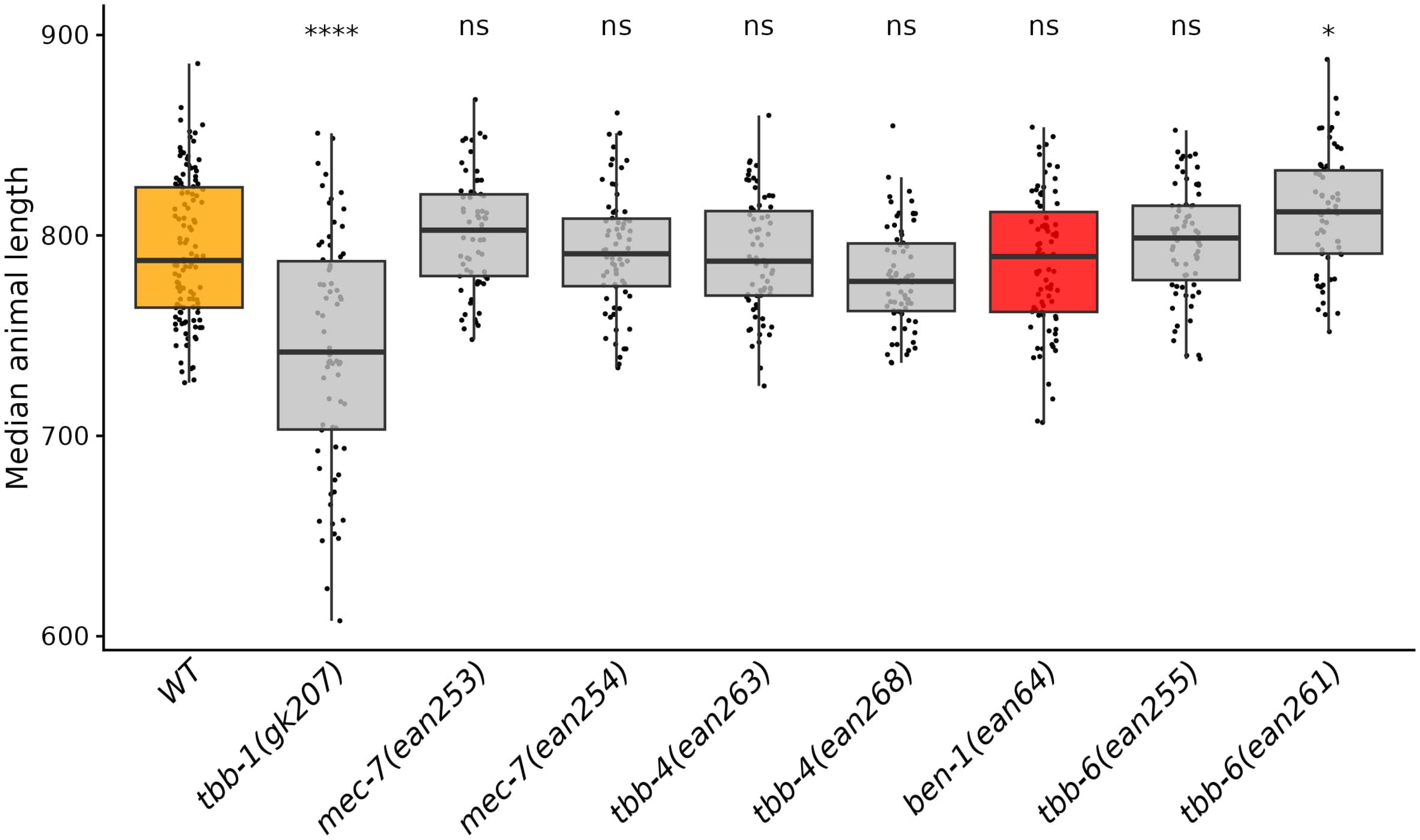
Loss of beta-tubulin genes affects animal lengths in control conditions. Median animal lengths of strains grown in 1% DMSO are shown. Each point represents the summarized measurements of an individual well containing five to 30 animals. Data are shown as box plots with the median as a solid horizontal line and the 75th and 25th quartiles on the top and bottom of the box, respectively. The top and bottom whiskers extend to the maximum point within the 1.5 interquartile range from the 75th and 25th quartiles, respectively. Statistical significance compared to the wild-type strain is shown above each strain (*p* < 0.05 = *, *p* < 0.0001 = ****, ANOVA with Tukey HSD).

**Supplemental Figure 7.**
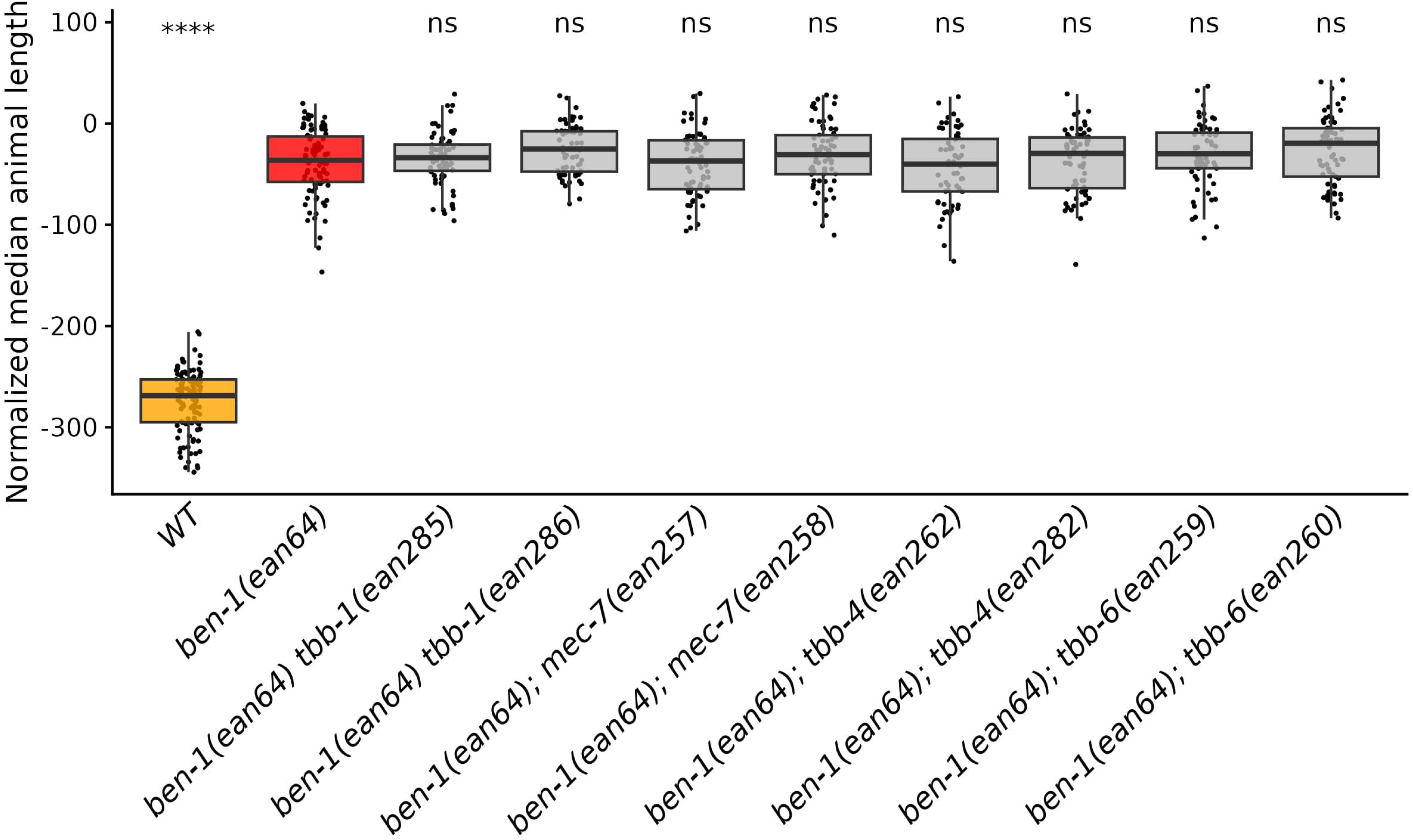
Additional loss of beta-tubulin genes in a Δ*ben-1* background did not confer a detectable level of increased ABZ resistance. Median animal lengths of strains grown in 30 µM ABZ that have been regressed for bleach effects and then normalized to the mean of all median animal lengths from the control condition are shown. Each point represents the summarized measurements of an individual well containing five to 30 animals. Data are shown as box plots with the median as a solid horizontal line and the 75th and 25th quartiles on the top and bottom of the box, respectively. The top and bottom whiskers extend to the maximum point within the 1.5 interquartile range from the 75th and 25th quartiles, respectively. Statistical significance compared to the Δ*ben-1* strain is shown above each strain (*p* < 0.05 = *, *p* < 0.0001 = ****, ANOVA with Tukey HSD).

**Supplemental Figure 8.**
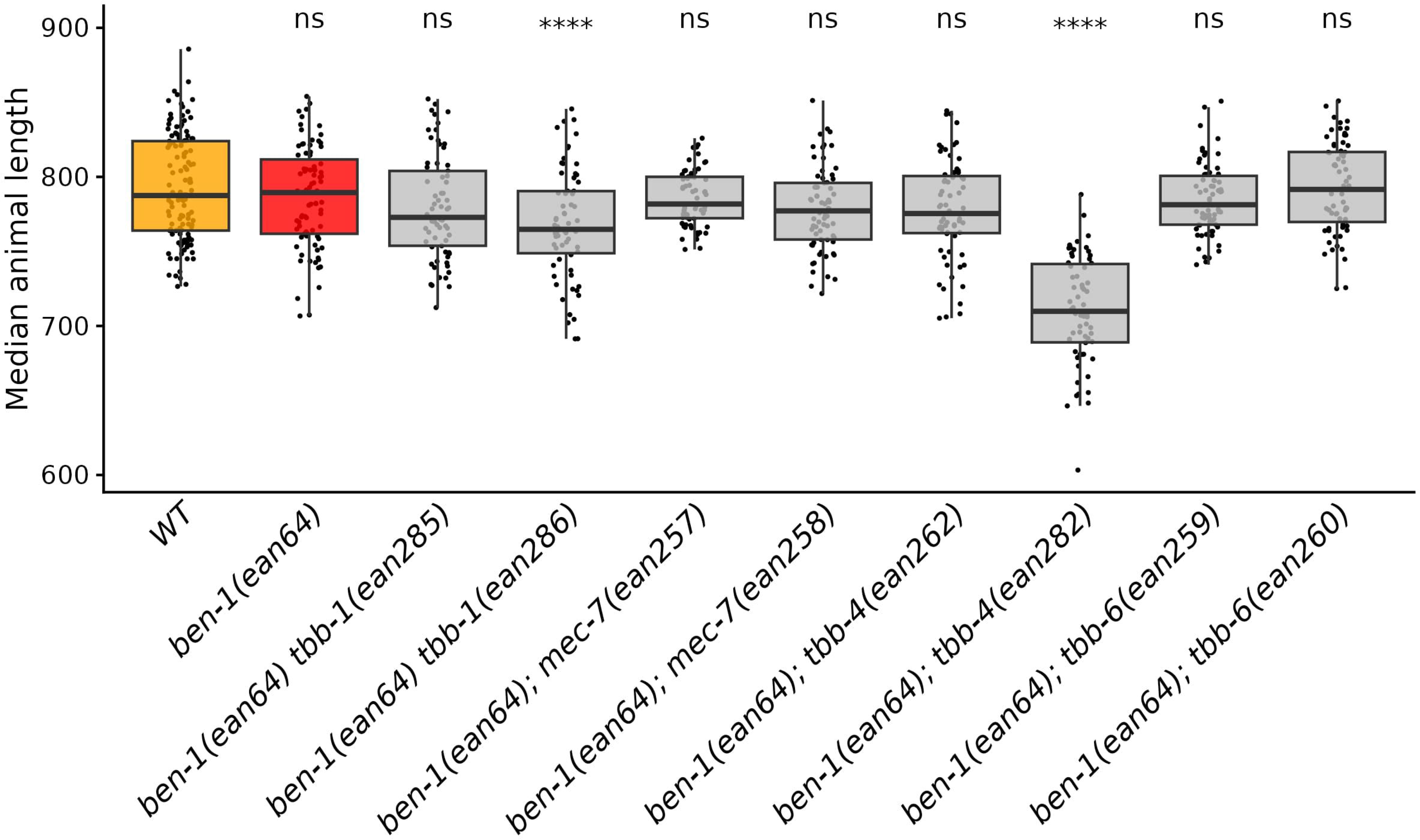
Loss of multiple beta-tubulins affects animal lengths in control conditions. Median animal lengths of strains grown in 1% DMSO are shown. Each point represents the summarized measurements of an individual well containing five to 30 animals. Where applicable, data from both independent edits in Assay 2 are shown. Data are shown as box plots with the median as a solid horizontal line, with the 75th and 25th quartiles on the top and bottom of the box, respectively. The top and bottom whiskers extend to the maximum point within the 1.5 interquartile range from the 75th and 25th quartiles, respectively. Statistical significance compared to the wild-type strain is shown above each strain (*p* < 0.05 = *, *p* < 0.001 = ***, *p* < 0.0001 = ****, ANOVA with Tukey HSD).

**Supplemental Figure 9.**
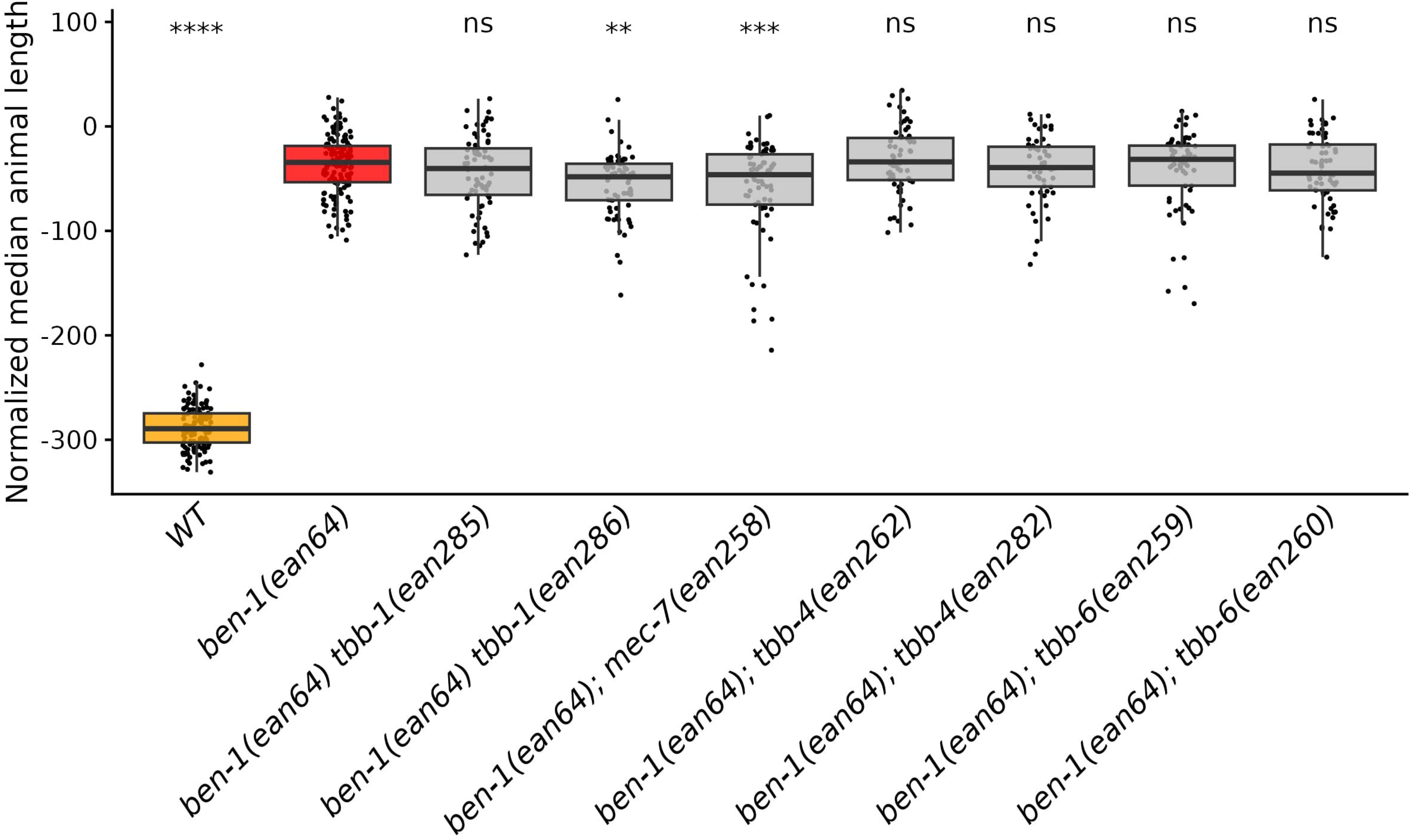
Additional loss of beta-tubulin genes in a Δ*ben-1* background did not confer a detectable level of increased ABZ resistance. Median animal lengths of strains grown in 30 µM ABZ that have been regressed for bleach effects and then normalized to the mean of all median animal lengths from the control condition are shown. Each point represents the summarized measurements of an individual well containing five to 30 animals. Data from both independent edits in Assay 1 are shown. Data are shown as box plots with the median as a solid horizontal line, with the 75th and 25th quartiles on the top and bottom of the box, respectively. The top and bottom whiskers extend to the maximum point within 1.5 interquartile range from the 75th and 25th quartiles, respectively. Statistical significance compared to the Δ*ben-1* strain is shown above each strain (*p* < 0.05 = *, *p* < 0.001 = ***, *p* < 0.0001 = ****, ANOVA with Tukey HSD).

**Supplemental Figure 10.**
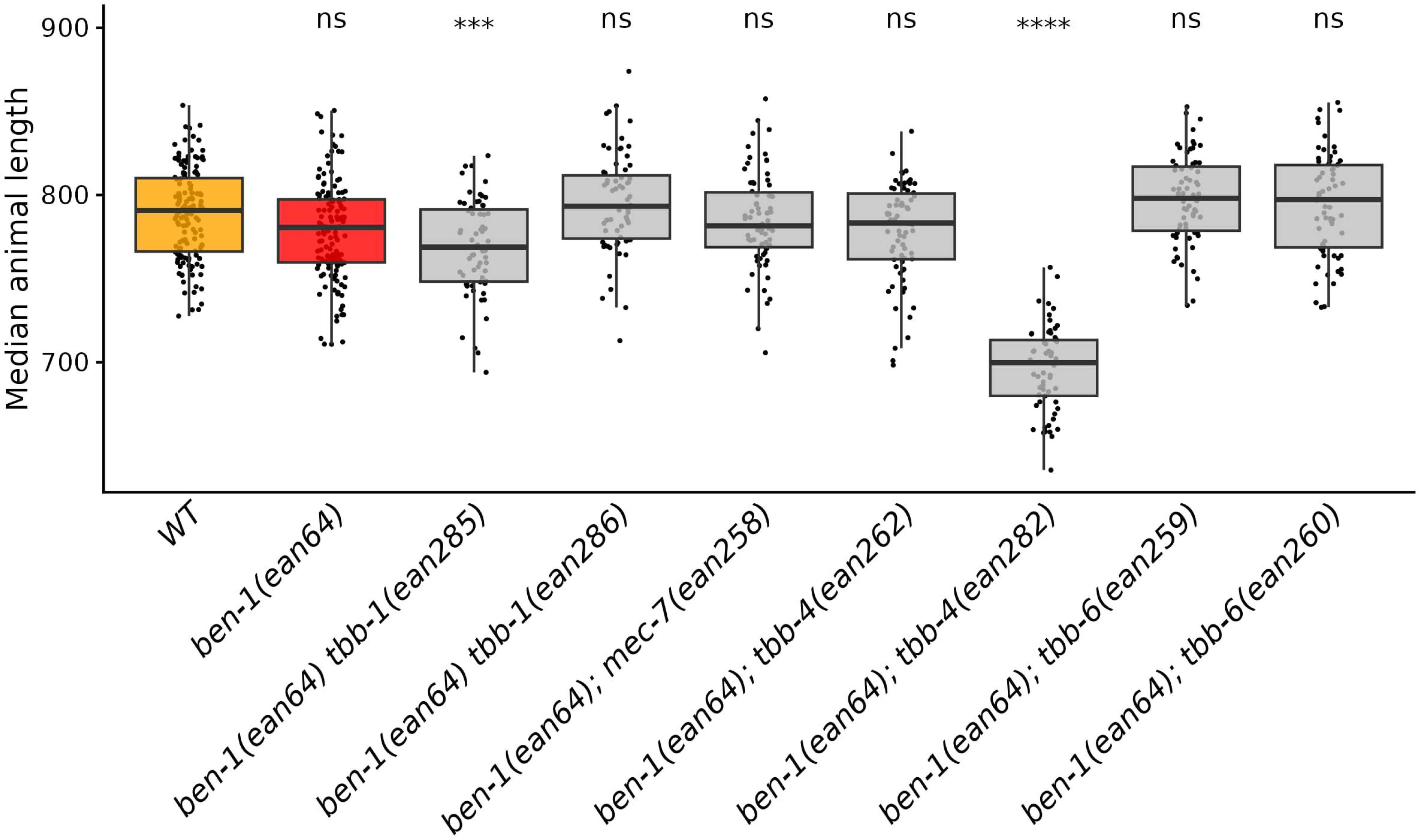
Loss of multiple beta-tubulin genes affects animal lengths in control conditions. Median animal lengths of strains grown in 1% DMSO are shown. Each point represents the summarized measurements of an individual well containing five to 30 animals. Data are shown as box plots with the median as a solid horizontal line and the 75th and 25th quartiles on the top and bottom of the box, respectively. The top and bottom whiskers extend to the maximum point within the 1.5 interquartile range from the 75th and 25th quartiles, respectively. Statistical significance compared to the wild-type strain is shown above each strain (*p* < 0.05 = *, *p* < 0.001 = ***, *p* < 0.0001 = ****, ANOVA with Tukey HSD).

## References

Andersen, E.C., Bloom, J.S., Gerke, J.P., Kruglyak, L., 2014. A variant in the neuropeptide receptor npr-1 is a major determinant of Caenorhabditis elegans growth and physiology. PLoS Genet. 10, e1004156.

Avramenko, R.W., Redman, E.M., Melville, L., Bartley, Y., Wit, J., Queiroz, C., Bartley, D.J., Gilleard, J.S., 2019. Deep amplicon sequencing as a powerful new tool to screen for sequence polymorphisms associated with anthelmintic resistance in parasitic nematode populations. Int. J. Parasitol. 49, 13–26.

Banerjee, S., Mukherjee, S., Nath, P., Mukherjee, A., Mukherjee, S., Ashok Kumar, S.K., De, S., Banerjee, S., 2023. A critical review of benzimidazole: Sky-high objectives towards the lead molecule to predict the future in medicinal chemistry. Results in Chemistry 6, 101013.

Boes, J., Eriksen, L., Nansen, P., 1998. Embryonation and infectivity of Ascaris suum eggs isolated from worms expelled by pigs treated with albendazole , pyrantel pamoate, ivermectin or piperazine dihydrochloride. Vet. Parasitol. 75, 181–190.

Boyd, W.A., Smith, M.V., Freedman, J.H., 2012. Caenorhabditis elegans as a Model in Developmental Toxicology, in: Harris, C., Hansen, J.M. (Eds.), Developmental Toxicology: Methods and Protocols. Humana Press, Totowa, NJ, pp. 15–24.

Chalfie, M., Thomson, J.N., 1982. Structural and functional diversity in the neuronal microtubules of Caenorhabditis elegans. J. Cell Biol. 93, 15–23.

Crombie, T.A., McKeown, R., Moya, N.D., Evans, K.S., Widmayer, S.J., LaGrassa, V., Roman, N., Tursunova, O., Zhang, G., Gibson, S.B., Buchanan, C.M., Roberto, N.M., Vieira, R., Tanny, R.E., Andersen, E.C., 2024. CaeNDR, the Caenorhabditis Natural Diversity Resource. Nucleic Acids Res. 52, D850–D858.

Dilks, C.M., Hahnel, S.R., Sheng, Q., Long, L., McGrath, P.T., Andersen, E.C., 2020. Quantitative benzimidazole resistance and fitness effects of parasitic nematode beta-tubulin alleles. Int. J. Parasitol. Drugs Drug Resist. 14, 28–36.

Dilks, C.M., Koury, E.J., Buchanan, C.M., Andersen, E.C., 2021. Newly identified parasitic nematode beta-tubulin alleles confer resistance to benzimidazoles. Int. J. Parasitol. Drugs Drug Resist. 17, 168–175.

Driscoll, M., Dean, E., Reilly, E., Bergholz, E., Chalfie, M., 1989. Genetic and molecular analysis of a Caenorhabditis elegans beta-tubulin that conveys benzimidazole sensitivity. J. Cell Biol. 109, 2993–3003.

Emms, D.M., Kelly, S., 2019. OrthoFinder: phylogenetic orthology inference for comparative genomics. Genome Biol. 20, 238.

Gibson, S.B., Ness-Cohn, E., Andersen, E.C., 2022. Benzimidazoles cause lethality by inhibiting the function of Caenorhabditis elegans neuronal beta-tubulin. Int. J. Parasitol. Drugs Drug Resist. 20, 89–96.

Hahnel, S.R., Zdraljevic, S., Rodriguez, B.C., Zhao, Y., McGrath, P.T., Andersen, E.C., 2018. Extreme allelic heterogeneity at a Caenorhabditis elegans beta-tubulin locus explains natural resistance to benzimidazoles. PLoS Pathog. 14, e1007226.

Hastie, A.C., Georgopoulos, S.G., 1971. Mutational resistance to fungitoxic benzimidazole derivatives in Aspergillus nidulans. J. Gen. Microbiol. 67, 371–373.

Howell, S.B., Burke, J.M., Miller, J.E., Terrill, T.H., Valencia, E., Williams, M.J., Williamson, L.H., Zajac, A.M., Kaplan, R.M., 2008. Prevalence of anthelmintic resistance on sheep and goat farms in the southeastern United States. J. Am. Vet. Med. Assoc. 233, 1913–1919.

Hurd, D.D., 2018. Tubulins in C. elegans. WormBook.

Kaplan, R.M., 2004. Drug resistance in nematodes of veterinary importance: a status report. Trends Parasitol. 20, 477–481.

Katoh, K., Misawa, K., Kuma, K.-I., Miyata, T., 2002. MAFFT: a novel method for rapid multiple sequence alignment based on fast Fourier transform. Nucleic Acids Res. 30, 3059–3066.

Kitchen, S., Ratnappan, R., Han, S., Leasure, C., Grill, E., Iqbal, Z., Granger, O., O’Halloran, D.M., Hawdon, J.M., 2019. Isolation and characterization of a naturally occurring multidrug-resistant strain of the canine hookworm, Ancylostoma caninum. Int. J. Parasitol. 49, 397–406.

Krücken, J., Fraundorfer, K., Mugisha, J.C., Ramünke, S., Sifft, K.C., Geus, D., Habarugira, F., Ndoli, J., Sendegeya, A., Mukampunga, C., Bayingana, C., Aebischer, T., Demeler, J., Gahutu, J.B., Mockenhaupt, F.P., von Samson-Himmelstjerna, G., 2017. Reduced efficacy of albendazole against Ascaris lumbricoides in Rwandan schoolchildren. Int. J. Parasitol. Drugs Drug Resist. 7, 262–271.

Kwa, M.S., Kooyman, F.N., Boersema, J.H., Roos, M.H., 1993. Effect of selection for benzimidazole resistance in Haemonchus contortus on beta-tubulin isotype 1 and isotype 2 genes. Biochem. Biophys. Res. Commun. 191, 413–419.

Kwa, M.S., Veenstra, J.G., Roos, M.H., 1994. Benzimidazole resistance in *Haemonchus contortus* is correlated with a conserved mutation at amino acid 200 in beta-tubulin isotype 1. Mol. Biochem. Parasitol. 63, 299–303.

Kwa, M.S., Veenstra, J.G., Van Dijk, M., Roos, M.H., 1995. Beta-tubulin genes from the parasitic nematode Haemonchus contortus modulate drug resistance in Caenorhabditis elegans. J. Mol. Biol. 246, 500–510.

Le, S.Q., Gascuel, O., 2008. An improved general amino acid replacement matrix. Mol. Biol. Evol. 25, 1307–1320.

Minh, B.Q., Nguyen, M.A.T., von Haeseler, A., 2013. Ultrafast approximation for phylogenetic bootstrap. Mol. Biol. Evol. 30, 1188–1195.

Minh, B.Q., Schmidt, H.A., Chernomor, O., Schrempf, D., Woodhams, M.D., von Haeseler, A., Lanfear, R., 2020. IQ-TREE 2: New Models and Efficient Methods for Phylogenetic Inference in the Genomic Era. Mol. Biol. Evol. 37, 1530–1534.

Mohammedsalih, K.M., Krücken, J., Khalafalla, A., Bashar, A., Juma, F.-R., Abakar, A., Abdalmalaik, A.A.H., Coles, G., von Samson-Himmelstjerna, G., 2020. New codon 198 β-tubulin polymorphisms in highly benzimidazole resistant Haemonchus contortus from goats in three different states in Sudan. Parasit. Vectors 13, 114.

Moya, N.D., Stevens, L., Miller, I.R., Sokol, C.E., Galindo, J.L., Bardas, A.D., Koh, E.S.H., Rozenich, J., Yeo, C., Xu, M., Andersen, E.C., 2023. Novel and improved Caenorhabditis briggsae gene models generated by community curation. bioRxiv. 10.1101/2023.05.16.541014

Nyaanga, J., Crombie, T.A., Widmayer, S.J., Andersen, E.C., 2021. easyXpress: An R package to analyze and visualize high-throughput C. elegans microscopy data generated using CellProfiler. PLoS One 16, e0252000.

Prichard, R.K., 1988. Anthelmintics and control. Vet. Parasitol. 27, 97–109.

R Core Team, 2020. R: A Language and Environment for Statistical Computing.

Salikin, N.H., Nappi, J., Majzoub, M.E., Egan, S., 2020. Combating Parasitic Nematode Infections, Newly Discovered Antinematode Compounds from Marine Epiphytic Bacteria. Microorganisms 8. 10.3390/microorganisms8121963

Saunders, G.I., Wasmuth, J.D., Beech, R., Laing, R., Hunt, M., Naghra, H., Cotton, J.A., Berriman, M., Britton, C., Gilleard, J.S., 2013. Characterization and comparative analysis of the complete Haemonchus contortus β-tubulin gene family and implications for benzimidazole resistance in strongylid nematodes. Int. J. Parasitol. 43, 465–475.

Shaver, A.O., Miller, I.R., Schaye, E.S., Moya, N.D., Collins, J.B., Wit, J., Blanco, A.H., Shao, F.M., Andersen, E.J., Khan, S.A., Paredes, G., Andersen, E.C., 2024. Quantifying the fitness effects of resistance alleles with and without anthelmintic selection pressure using Caenorhabditis elegans. bioRxiv. 10.1101/2024.02.01.578300

Shaver, A.O., Wit, J., Dilks, C.M., Crombie, T.A., Li, H., Aroian, R.V., Andersen, E.C., 2023. Variation in anthelmintic responses are driven by genetic differences among diverse C. elegans wild strains. PLoS Pathog. 19, e1011285.

Shaver, A.O., Wit, J., Dilks, C.M., Crombie, T.A., Li, H., Aroian, R.V., Andersen, E.C., 2022. Variation in anthelmintic responses are driven by genetic differences among diverse C. elegans wild strains. bioRxiv. 10.1101/2022.11.26.518036

Sheir-Neiss, G., Lai, M.H., Morris, N.R., 1978. Identification of a gene for beta-tubulin in Aspergillus nidulans. Cell 15, 639–647.

Silvestre, A., Cabaret, J., Humbert, J.F., 2001. Effect of benzimidazole under-dosing on the resistant allele frequency in Teladorsagia circumcincta (Nematoda). Parasitology 123, 103–111.

Venkatesan, A., Jimenez Castro, P.D., Morosetti, A., Horvath, H., Chen, R., Redman, E., Dunn, K., Collins, J.B., Fraser, J.S., Andersen, E.C., Kaplan, R.M., Gilleard, J.S., 2023. Molecular evidence of widespread benzimidazole drug resistance in Ancylostoma caninum from domestic dogs throughout the USA and discovery of a novel β-tubulin benzimidazole resistance mutation. PLoS Pathog. 19, e1011146.

Widmayer, S.J., Crombie, T.A., Nyaanga, J.N., Evans, K.S., Andersen, E.C., 2022. C. elegans toxicant responses vary among genetically diverse individuals. Toxicology 479, 153292.

Wit, J., Dilks, C.M., Andersen, E.C., 2021. Complementary Approaches with Free-living and Parasitic Nematodes to Understanding Anthelmintic Resistance. Trends Parasitol. 37, 240–250.

Yang, Z., 1994. Maximum likelihood phylogenetic estimation from DNA sequences with variable rates over sites: approximate methods. J. Mol. Evol. 39, 306–314.

Zamanian, M., Cook, D.E., Zdraljevic, S., Brady, S.C., Lee, D., Lee, J., Andersen, E.C., 2018. Discovery of genomic intervals that underlie nematode responses to benzimidazoles. PLoS Negl. Trop. Dis. 12, e0006368.

Zhang, G., Roberto, N.M., Lee, D., Hahnel, S.R., Andersen, E.C., 2022. The impact of species-wide gene expression variation on Caenorhabditis elegans complex traits. Nat. Commun. 13, 3462.

